# Inhibition of the Lysosomal Amino Acid Sensor SLC38A9 by the Membrane Microprotein SPAR

**DOI:** 10.64898/2026.08.07.743590

**Authors:** Ampon Saeher, Xuelang Mu, Tamir Gonen

## Abstract

Long noncoding RNAs encode for microproteins that regulate cellular functions. Small regulatory peptide of amino acid response (SPAR) is a microprotein in the lysosome that responds to amino acid availability of the cell. In this study, we investigated the interactions between SPAR and SLC38A9, a lysosomal amino acid transporter and receptor involved in the mechanistic target of rapamycin 1 (mTORC1) pathway. We found that SPAR binds SLC38A9 and inhibits arginine transport in SLC38A9. Moreover, the downstream recruitment of Rag GTPases is also inhibited when SPAR is present in SLC38A9 liposomes. Docking model shows potential interactions between SPAR and SLC38A9. Together, these findings reveal the mechanism of mTORC1 inhibition through microprotein SPAR and illustrates the power of non long coding RNAs in altering cellular functions.

**Statement of Significance:** Microproteins encoded from long noncoding RNAs are emerging as critical regulators of many pathways. This study investigates a novel mechanism of SPAR microprotein that directly regulates the mechanistic target of rapamycin complex1 (mTORC1) signaling pathway through the lysosomal amino acid transporter SLC38A9. SPAR blocks both arginine transport and the downstream recruitment of Rag GTPases. These findings provide critical results in how SPAR controls cellular amino acid availability, while broadly highlighting the powerful regulatory mechanism of microproteins in cellular processes.

## Introduction

Microproteins are generated from small open reading frames which produce protein sequences that are less than 100 amino acids in length (1) . These proteins have been shown to play a broad range of roles in cellular functions including cell development, differentiation, stress signaling and metabolism (2). For instance, a 34 amino acid long noncoding RNAs (lncRNA) has been discovered to regulate the function of a SERCA, a calcium ATPase and regulate muscle function by displacing SERCA inhibitors including phospholamban, sarcolipin, and myoregulin (3). Therefore, there is a growing interest in investigating the functions of microproteins.

SPAR (small regulatory polypeptide of amino acid response or LINC00961) is a 90 amino acid polypeptide that regulates mTORC1 (mechanistic target of rapamycin complex 1) localization on the lysosome (4). The mTORC1 is a multiprotein signaling kinase that regulates growth, protein synthesis, and metabolism by using the environmental cues available in cells such as amino acids availability. Upstream regulators of mTORC1 including Rag GTPases, SPAR interacts with v-ATPase and Ragulator/Rag GTPase complexes to regulate mTORC1 function (4). In the presence of SPAR, mTORC1 complex is inactivated and cells become irresponsive to amino acid stimulation or starvation. Therefore, SPAR microprotein has been proposed as an inhibitor of the mTORC1 pathway (4). In the absence of SPAR, the mTORC1 pathway is hyperactivated and this leads to cell proliferation and stem cell differentiation. SPAR has been shown to interact with the lysosomal V-ATPase to negatively regulate mTORC1 pathway (4) . However, the relationship between SPAR and the intricate mechanism of amino acid sensing in the mTORC1 pathway remains unclear. In particular, how SPAR interacts with the lysosomal protein SLC38A9 is unknown. SLC38A9 functions as a transceptor, transporter and receptor that regulates mTORC1. We hypothesized the interactions between SPAR and SLC38A9 tuned the lysosomal amino acid availability and the activation of mTORC1.

To investigate potential interactions between SPAR and SLC38A9 we used binding experiments, pull-down assays, radiolabeled substrate uptake assays, and computational docking experiments. We discovered that SPAR directly binds SLC38A9 and modulates its activity. We found that the presence of SPAR inhibits SLC38A9 Arginine uptake activity. Moreover, liposome pull-down assays of SLC38A9 and Rag GTPases showed that downstream signaling of mTORC1 was also inhibited in the presence of SPAR. This interaction was probed using a three-dimensional structural model and site directed mutagenesis experiments which confirmed our findings.

## Materials and Methods

### Protein expression

DNA sequences of zebrafish and human SLC38A9 were cloned into *pF*astbac1 vectors for expression in *Spodoptera frugiperda* sf-9 cells using the baculovirus technique as previously described (6). DNA sequences for human and mouse SPAR (GenBank: AL135841, AL773539 respectively) were cloned into *p*mal-c2x vectors and expressed with maltose binding protein as a fusion protein for protein solubility. These were then transformed into BL21-Gold competent cells using carbenicillin plates for positive colony selection. Overnight growth was done at 37°C in 100mL of Luria-Bertani media (LB) supplemented with carbenicillin. This culture was transferred to large scale growth the next day, grew till the OD600 of 1.0, and induced with 0.5mM IPTG.

### Site Directed Mutagenesis

Site-directed mutagenesis was performed using the QuickChange Site-Directed Mutagenesis Kit (Agilent) using primers listed in SI table 2. The correct DNA plasmid sequences were confirmed by sequencing.

### Generation of SLC38A9 variants

The sequence encoding hSLC38A9 (Genbank Q8NBW4) was cloned into the *p*Fastbac 1 vector (Invitrogen) with a sequence encoding a C-terminal 8X His tag and a thrombin cleavage site. The SPAR fused-SLC38A9 (SS-fusion) protein sequence was cloned to have an N-terminal 8X His tag and a thrombin cleavage site before the drSLC38A9 sequence, followed by a 6X Gly-link and mSPAR (Genbank AL773539) sequence and a flag tag. All wild-type and variants of SLC38A9 were confirmed by whole-plasmid sequencing. Verified plasmids were transformed into DH10Bac cells (From Thermo Fisher Scientific MAX Efficiency DH10Bac competent cells) for preparation of bacmids. Recombinant baculovirus was generated and used for transfection following the protocol for the Bac-to-Bac Baculovirus Expression System (Invitrogen). Wild-type SLC38A9 and SPAR fused-SLC38A9 proteins were overexpressed in *Spodoptera frugiperda* sf-9 insect cells, which were harvested at 60 hours after P2 virus infection. The *Spodoptera frugiperda* sf-9 insect cells are from Expression Systems, adapted in ESF 921.

### SLC38A9 Protein and Proteoliposomes Preparation

The protocol for purifying SLC38A9 wild-type protein and SS-fusion protein was described by (7). The purified membrane proteins were in a buffer containing 20mM Tris 8.0, 150mM NaCl, and 0.2% DM, for proteoliposome reconstitution and following assays.

Chloroform-dissolved chicken egg phosphatidylcholine (egg-PC, Avanti Polar Lipids) was evaporated using dry nitrogen to yield a lipid film in a small glass vial and further dried under vacuum overnight. The lipids were hydrated in the inside buffer (20 mM Tris 8.0, 90 mM KCl, 10 mM NaCl) at 25 mg/mL by vortexing for 3 minutes and then aged at room temperature for 1 hour. Liposomes were clarified by 5 rounds of freezing and thawing in liquid nitrogen and extruded through a 100 nm membrane with 21 passes (Millipore). The liposomes were pre-incubated with 1% n-octyl-b-D-glucoside (β-OG) and 1 mM DDT for 1 hour at 4°C before protein reconstitution. Purified wild-type SLC38A9 and purified SPAR proteins were incorporated at a 1:50 (w/w) ratio into destabilized liposomes for 1 hour in the 4°C rotator. Glycerol-supplemented protein buffer was used in lieu of the SLC38A9 protein in liposome-only control groups. The detergents were removed by incubation overnight with 200mg per reaction Bio-Beads, and the proteoliposomes were further incubated with 40mg per reaction fresh Bio-Beads for an additional hour. The proteoliposomes and liposome-only controls were collected using an ultracentrifuge at 100,000 x *g* for 30 minutes at 4°C and then resuspended in an outside buffer (100 mM NaCl and 20mM Mes 6.0) to a final lipid concentration of 40 μg/μL.

### Transport Assay

Transport reactions were initiated by adding 0.5 μM L-[^3^H]-arginine (American Radiolabeled Chemicals, Inc, batch/Lot #220722) to 50 μL of proteoliposomes. Assays of Liposome-only controls were carried out in parallel to experimental groups as negative controls. All buffers were chilled, and assays were performed at a 30°C metal bath. At the 10-minute time points, the uptake reaction was saturated so that proteoliposomes were filtered, washed by 10mL of ice-cold wash buffer (outside buffer with 10 mM unlabeled L-arginine), and collected on 0.22 μm nitrocellulose membranes (Millipore) which had been pre-wet by washing buffer. After washing, each membrane was dried under vacuum for exactly 1 minute and transferred into a glass vial with 10 mL scintillation fluid for counting. Non-specific adsorptions of L-[^3^H]-arginine by liposomes-only controls were subtracted from experimental measurements. All experimental and control groups were repeated three times.

### hRags Protein Preparation

RagA/C GTPases were expressed as previously published (12, 13). Namely, RagA/C GTPases (cloned in PetDuet-1 vector) were transformed into BL21(DE3) cells. Cells were grown in LB media at 37C until OD600 reached 1.5 and 0.4mM IPTG was added to the media to induce expression. Cells were then grown overnight at 24C. Cells overexpressing RagA/C GTPases were centrifuged at 5000g for 10 mins and collected for protein purification. Cells were resuspended in 20mM Tris, 150mM NaCl pH 8.0 buffer with protease inhibitors and homogenized and lyzed under a high pressure in the microfluidizer for 3 cycles. Cell debris was centrifuged at 10000g for 30 minutes, then the supernatant was applied to cobalt resin and incubated for 1 hour with stirring.

Then, resin was washed with 20mM Tris 8.0, 150mM NaCl, 20mM imidazole buffer for 10x column volume. Thrombin (1U/μL) was added to cleave the proteins on column overnight in 4°C. Cleaved protein was further purified in size exclusion chromatography and fractions with the highest amount of protein were pooled and flash-frozen with liquid nitrogen for further use.

### Pull-Down Assay and Western Blotting

Purified drSLC38A9 and SPAR fused-SLC38A9 (SS-Fusion) proteins were reconstituted into liposomes at a protein-to-lipid ratio of 1:50 (w/w). Glycerol-supplemented protein buffer was used in lieu of SLC38A9 or SS-fusion proteins in the liposome-only control group. The amount of protein incorporated into proteoliposomes was quantified by SDS-PAGE using BSA standards (Thermo Scientific) run on the same gel. Purified Rag GTPases were prepared as described above. Rag proteins were first activated by incubation with a 10-fold molar excess of GDP for 30 minutes at room temperature, followed by the addition of MgCl_2_ to a final concentration of 20mM. To assemble the SLC38A9-Rag and SPAR-fused SLC38A9-Rag complexes, proteoliposomes were incubated with activated Rag Proteins at a 1:4 molar ratio (mol/mol) in the presence of 20 μM arginine overnight at 4°C. The following day, proteoliposomes and control groups were pelleted by ultracentrifugation at 100,000 x *g* for 30 minutes at 4°C. Supernatants were collected to assess unbound proteins by SDS-PAGE. Pellets were washed with pre-chilled arginine-containing outside buffer, resuspended in the same buffer, and pelleted again under the same conditions. This washing step was repeated three times in total. Supernatant and pellet fractions from the final wash were collected and analyzed by SDS-PAGE and Western Blot.

Proteins were then transferred onto PVDF membranes using a semi-dry transfer method. Membranes were blocked with 5% milk in PBST buffer and incubated overnight at 4°C with primary antibodies against SLC38A9 (HPA0437785, Sigma), and Rag A (4357, Cell Signaling) with Rag C (5466, Cell Signaling). Membranes were subsequently incubated with a donkey anti-rabbit secondary antibody (SA1-200, Invitrogen) for 1 hour at room temperature. Signals were detected using an ECL detection kit (Bio-Rad).

### Microscale thermophoresis (MST) experiments

Purified SPAR protein was labeled with the NHS-RED protocol published on the NanoTemper website. The degree of labeling was measured to be ∼1 before proceeding with the binding experiments. Labeled protein (either SPAR or SLC38A9) was diluted to ∼2μM and diluted to 40nM for the MST experiments. Serial dilutions of unlabeled proteins were made for 16 samples starting from 200μM. Then labeled proteins (20μM) were added to each serially diluted sample. Samples were mixed and centrifuged to remove aggregations. These samples were then loaded into the capillary tubes for fluorescence measurement on the NanoTemper MST machine. New samples were repeated at least 3 times for each experiment.

### Docking experiments

Protein-protein docking was performed using the ClusPro web server. The input models used were from the previously published structure of SLC38A9 (PDB: 6C08) (5) and the alphafold model of SPAR protein. The docking was executed using the PIPER docking program which employs a Fast Fourier Transform (FFT) based rigid body search with pairwise interaction potentials. The lowest energy conformations generated by PIPER were clustered using an algorithm based on root-mean-square deviation (RMSD). The centers of the most populated clusters were selected as representative models of the complex. The most probable docking model was then validated with binding experiments.

## Results

### SPAR binds SLC38A9

SLC38A9 and SPAR proteins were expressed and purified as described. Microscale thermophoresis (MST) assays are performed on SPAR and SLC38A9 to determine the binding affinity of the complex. Due to the aggregation in SPAR protein, we designed a maltose binding protein (MBP) fusion construct with SPAR that is overexpressed in high yield and purified as a soluble protein which is used throughout this study. SPAR was labeled with a Lys-labeling kit as described in the methods section. Labeled SPAR was used for serial dilution of SLC38A9 and the change in fluorescence intensity was recorded on the MST machine. We performed these MST binding experiments on zebrafish SLC38A9 (drSLC38A9), human SLC38A9 (hSLC38A9), mouse SPAR (mSPAR) and human SPAR (hSPAR) proteins. The binding curves were then fitted to obtain dissociation constants (Kd) of 2.6 μM (Figure 1a) between drSLC38A9 and mSPAR and 4.7μM between hSLC38A9 and hSPAR. Truncation of the SPAR C terminal (SPARΔC) makes the interactions between SLC38A9 and SPAR weaker (Kd of 4.7μM, supplemental Figure 1). These results show that SPAR interacts with SLC38A9 *in vitro* and suggest that the C terminal of SPAR might play a role in the interactions between the two proteins.

**Figure 1.**
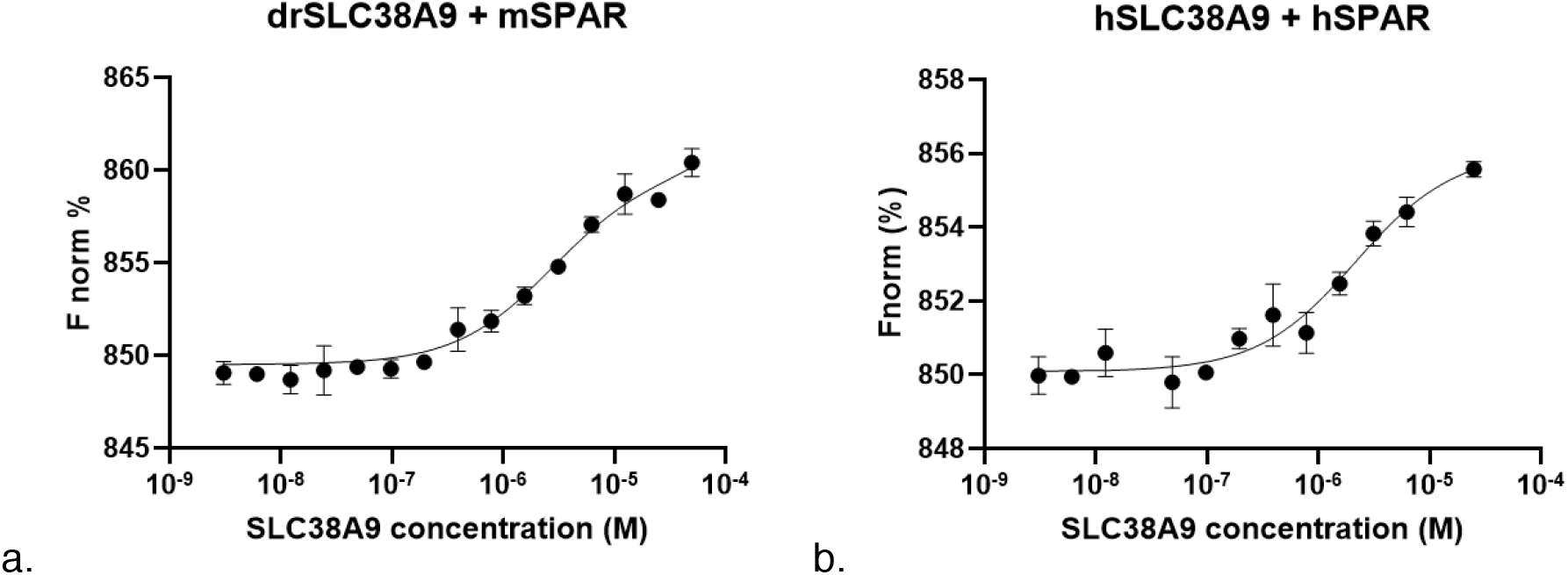
Microscale Thermophoresis (MST) Experiments of SLC38A9 and SPAR proteins show that SPAR binds to SLC38A9. (a) MST binding experiment of zebrafish SLC38A9 (drSLC38A9) and mouse SPAR (mSPAR). (b) MST binding experiment of human SLC38A9 (hSLC38A9) and human SPAR (hSPAR).

### SPAR inhibits Arginine uptake function in SLC38A9

Arginine uptake assays have been performed on SLC38A9 to show the intrinsic activity of the SLC38A9 transporter in liposomes (5,6) Previously, we have shown that SLC38A9 transports arginine in a sodium-dependent and a pH-regulated manner (7). To investigate how SPAR affects the arginine transport activity of SLC38A9, we reconstituted SLC38A9 and both SPAR and SLC38A9 into liposomes and performed scintillation counting of ^3^H-Arginine uptake (Figure 2A). The results showed that SPAR inhibits Arginine uptake in SLC38A9 to a similar level as the negative control SPAR and the empty liposomes without SLC38A9 (Figure 2B). We also performed Arginine uptake assays of the SLC38A9-SPAR fusion protein (SS-fusion) and observed a significant reduction of Arginine uptake compared to wild type drSLC38A9 (Figure 2C). Similarly, the SS-fusion proteoliposome showed a similar level of ^3^H-arginine uptake as the control empty liposomes (Figure 2C). Since SPAR is expressed as a fusion protein with a maltose binding protein (MBP), we designed a hSPAR construct that has a cleavable MBP to ensure that MBP does not affect the ^3^H-Arginine uptake activity of SLC38A9. Arginine uptake assays were performed on co-reconstituted MBP-free hSPAR and hSLC38A9, MPB-free hSPAR inhibits arginine uptake activity similar to the MBP fusion SPAR (supplemental figure 2). Full length hSLC38A9 (FL-hSLC38A9) and N-terminal truncated hSLC38A9 (ΔN-hSLC38A9) both showed similar levels of arginine uptake into the liposomes. However, hSPAR exhibits ∼1.6 fold higher inhibition of arginine uptake in ΔN-hSLC38A9 than FL-hSLC38A9. This suggests that the N-terminal of SLC38A9 might play a role in reducing the interactions with SPAR thus reducing its inhibition in the arginine uptake assay.

**Figure 2.**
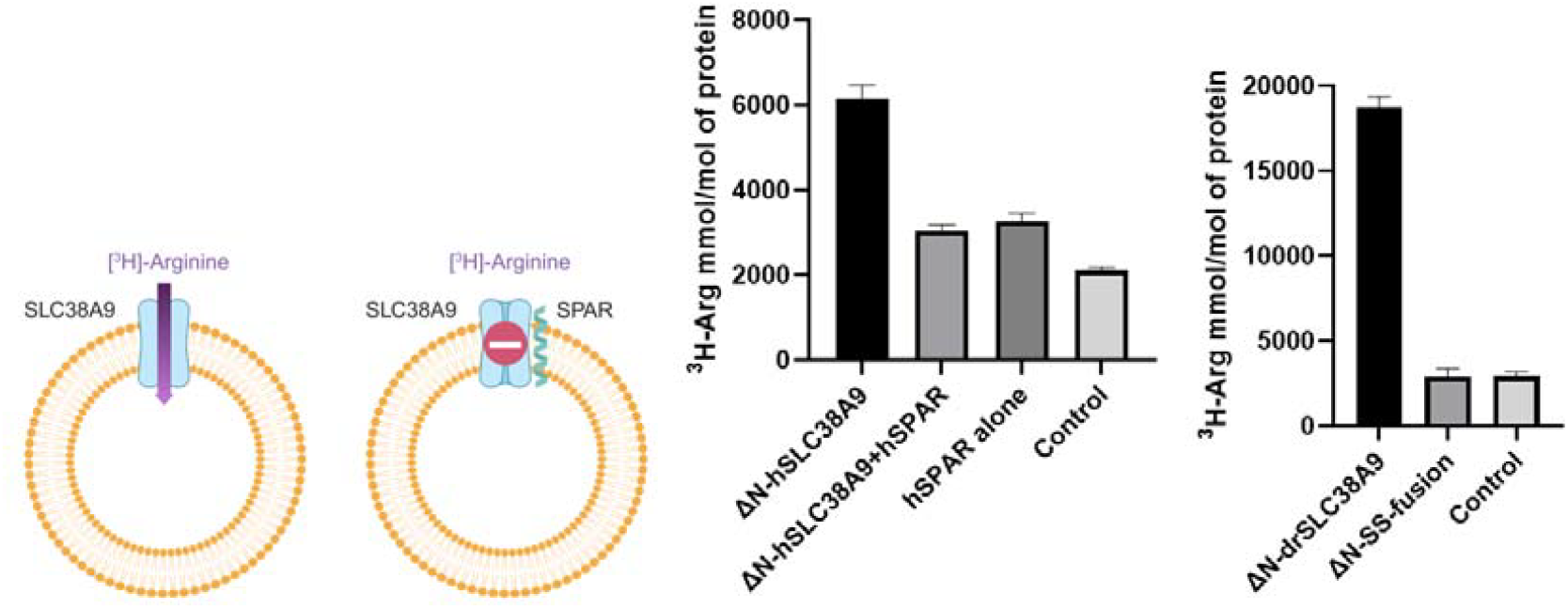
SPAR inhibits arginine uptake in SLC38A9. (a) Schematic of the liposome uptake experiments. (b) ^3^H-Arginine uptake experiments of SLC38A9 and SPAR. (c) ^3^H-Arginine uptake experiments of SLC38A9 and SS-fusion

**Figure 3.**
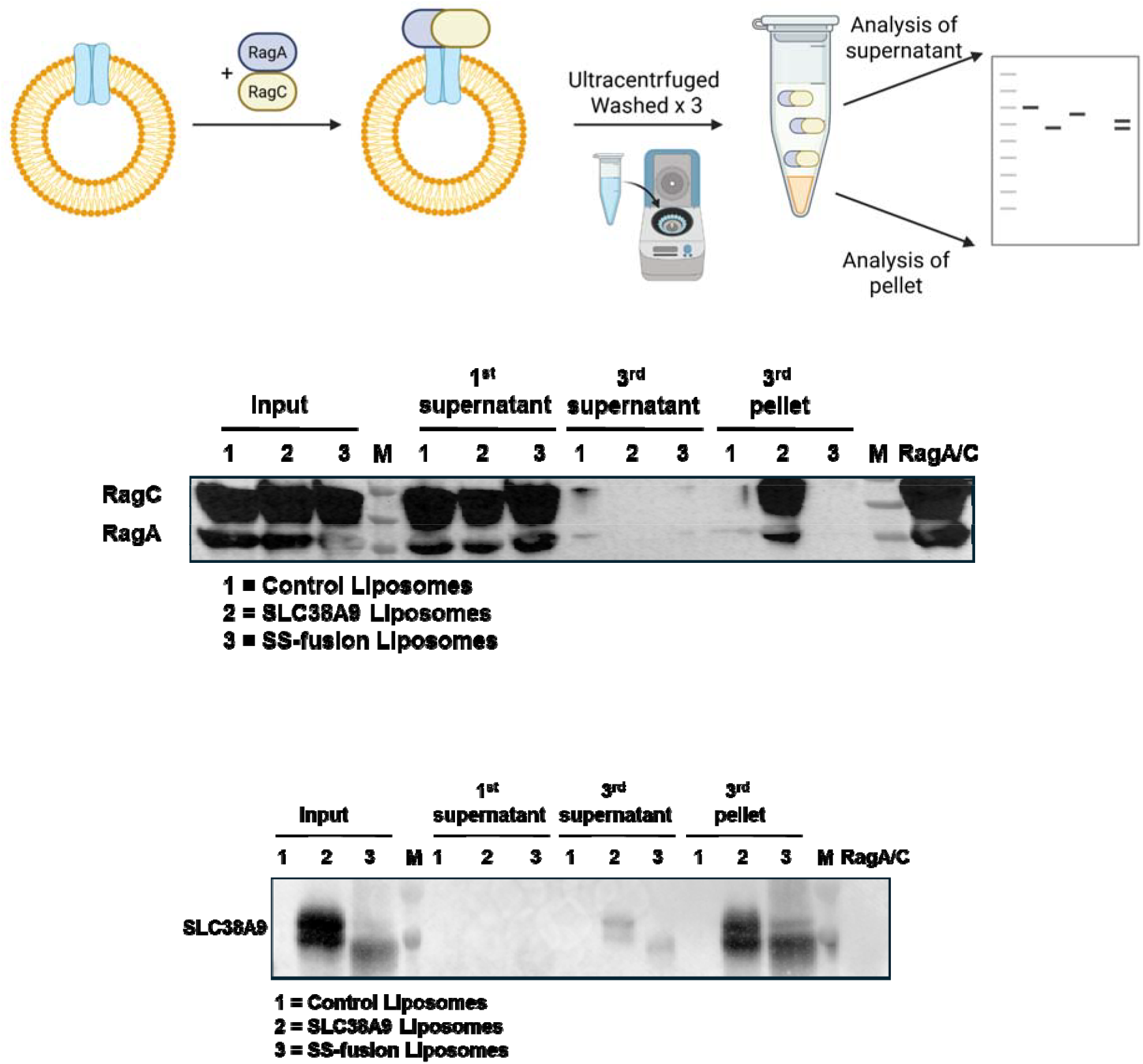
(a) Schematic of liposome pull-down assays. SLC38A9 or SLC38A9 fused with SPAR protein (SS-fusion) were reconstituted in liposomes and RagA/C GTPases were bound overnight. The proteoliposome mixtures were then ultracentrifuged and washed 3 times with a wash buffer. Samples of supernatant and pellet were taken for western blot analysis. (b) Anti-RagA/C western blot analysis of proteoliposome samples of empty liposome as control (1), SLC38A9 (2), and SS-fusion (3). Input proteoliposomes were sampled before ultracentrifugation. First supernatant samples were taken from the first ultracentrifugation step. Third supernatant and pellet samples were taken from the 3rd wash step. Protein marker (M) shows molecular weight of 55, 40, 35 kDa from top to bottom. (c) Anti-SLC38A9 western blot of the same samples in Figure 3b. Protein marker (M) shows molecular weight of 55and 40 kDa from top to bottom.

**Figure 4.**
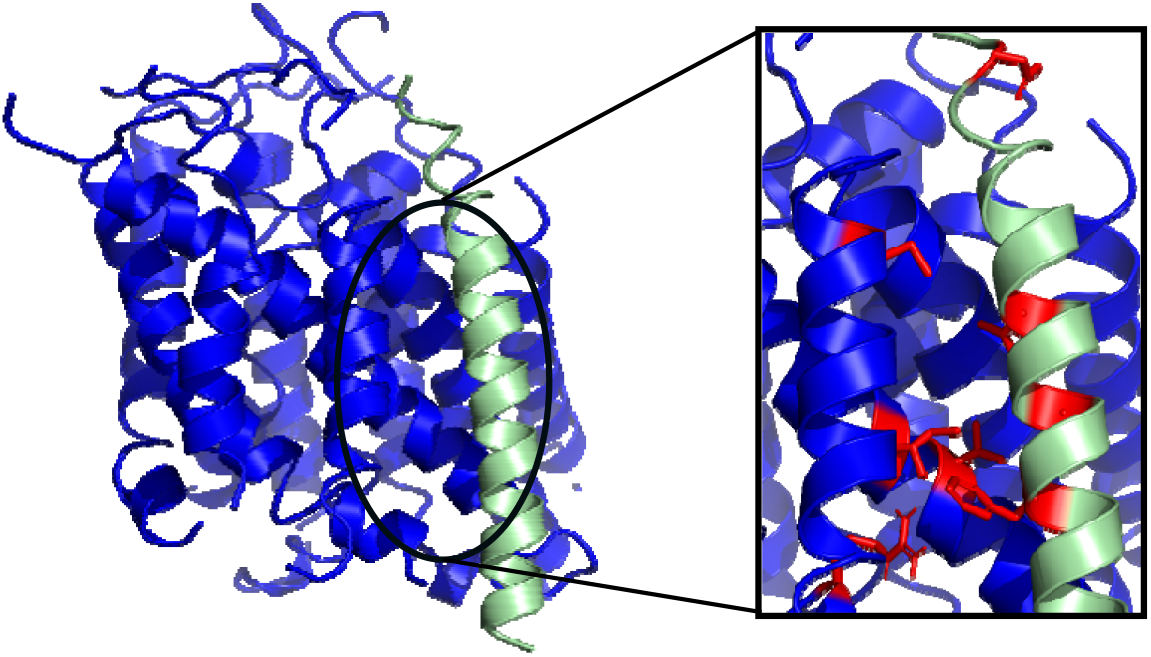
Docking experiment of SPAR and SLC38A9 via Cluspro (9, 10). Residues highlighted in red are with the vicinity of the protein-protein interface. Residues on SPAR are T13, T16, T20, and D29. Residues on SLC38A9 are V195, I198, M483, and R541.

### SPAR inhibits Rag GTPase binding in the liposomal pull-down assays

To test if the binding between SLC38A9 and Rag GTPases is also disrupted by SPAR *in vitro*, we performed pull-down assays in liposomes. Since SPAR is predicted to have a single helix and a long c tail in Alphafold and its topology *in vivo* has been previously established (4), we designed a fusion construct where drSLC38A9 is fused with SPAR protein on the C-terminal (SS-fusion). SLC38A9 and SLC38A9 with SPAR protein (SS-fusion) were reconstituted and Rag GTPases were added to the proteoliposomes. These proteoliposomes were incubated overnight with RagA/C GTPases and 200μM Arginine. Then, the proteoliposomes were ultracentrifuged and washed twice with a buffer. The components in proteoliposome and the supernatant fractions were then analyzed by western blots (Figure 2a). Three proteoliposome samples were the negative control empty liposomes (Figure 2b, sample 1), SLC38A9 proteoliposomes (Figure 2b, sample 2), and SS-fusion proteoliposomes (Figure 2b, sample 3). The first supernatant showed excess RagA/C GTPases being washed off of the proteoliposome samples (Figure 2b, 1st supernatant). The wash steps were completed after the third wash cycle as there were no more RagA/C GTPases in the supernatant (Figure 2b, 3rd supernatant). SLC38A9 pulled down RagA/C GTPases as there are bands that correlate to the molecular weights of RagA/C GTPases in the 3rd pellet fraction of SLC38A9 (Figure 2b, 3rd pellet, sample 2). The fusion protein (SS-Fusion) did not pull down RagA/C GTPases and showed no bands that correlate to RagA/C GTPase in the 3rd pellet fraction (Figure 2b, 3rd pellet, sample 3). The western blot analysis shows that the supernatant from the proteoliposomes containing SLC38A9 does not have the bands correlates to RagA/C GTPases but the supernatants from the proteoliposomes containing SS-Fusion have them. These results show that the interactions between SLC38A9 and SPAR inhibit the binding to RagA/C GTPases. Full SDS-PAGE gels and western blot membranes are shown in Supplemental Figure 3.

### Docking experiments highlight interactions between SLC38A9 and SPAR

To understand the atomic interactions between SLC38A9 and SPAR, we predicted the structure of SPAR for docking experiments using alphafold (8) and performed docking experiments between the experimental structure of SLC38A9 (5) and SPAR using cluspro (9, 10). Docking analysis using the structure of SLC38A9 (PDB: 6C08) and the alphafold predicted structure of SPAR showed that the single helix of SPAR interacted with the transmembrane domains 2, 9, and 10 of SLC38A9. To validate the docking results, we mutated several amino acid residues on SPAR that are within the vicinity of the protein-protein interaction interface (i.e., less than 5 Å) including F13, T16, T20, C21, and D29. These residues were mutated to alanine and MST experiments were performed to investigate if they still bind to SLC38A9. All these SPAR mutants abolish binding to SLC38A9 (supplemental figure 4). In addition, we mutated residues on SLC38A9 that are within the vicinity of the protein-protein interaction interface to alanine including V195, I198, M483, and R541. We then performed MST experiments and observed that the binding to SPAR is abolished in these mutants (supplemental figure 5).

## Discussion

Regulation of mTORC1 activity is important in cell signaling and survival. SLC38A9 has been shown to regulate mTORC1 pathway through its nutrient sensing function (11). The structure of SLC38A9 has 11 transmembrane domains with a pseudo two-fold symmetry with one transmembrane domain flanking one side of the transporter (5). This study proposes that a single transmembrane domain microprotein called SPAR regulates the function of SLC38A9. As shown in a previous study (4), SPAR negatively regulates the mTORC1 pathway by amino acid stimulation and interacts with v-ATPase. Since SLC38A9 is the transceptor responding to the amino acid concentrations in the mTORC1 pathway, we showed in this study that SPAR also interacts with SLC38A9.

Here we report the data on binding assays of SPAR and SLC38A9, inhibition of ^3^H-Arginine uptake assays of SLC38A9 by SPAR, inhibition of the pull-down assays of RagA/C GTPases by SPAR, and a docking model of SPAR and SLC38A9. SPAR binds and interacts with SLC38A9 both in detergent and liposome. It also inhibits the amino acid transport function intrinsic to SLC38A9. Moreover, downstream recruitments of RagA/C GTPases are also inhibited by SPAR. This means that the downstream mTORC1 activation is also inhibited in agreement with the previous findings that SPAR regulates the mTORC1 activation (4). Thus, SPAR regulates the mTORC1 pathway in several ways including the interaction with v-ATPase and SLC38A9. Further structural studies are needed in the future to understand the interactions between these protein complexes and their relations to mTORC1. Importantly, since SPAR is relatively small compared to other membrane proteins involved in the mTORC1 pathway, it can also be a good therapeutic candidate for cancer inactivation. This study further emphasizes the significance of small long non coding RNAs (lncRNAs) and their interactions with protein complexes specifically involved in the mTORC1 inactivation. Future structural studies are needed to illustrate how this interaction affects the mTORC1 pathway and downstream phosphorylation.

## Abbreviations

mTORC1: mechanistic target of rapamycin complex1
lncRNAs: long non coding RNAs
SPAR: Small regulatory peptide of amino acid response
SLC38A9: Solute Carrier Family 38 Member 9
MST: Microscale Thermophoresis
SS-Fusion: SPAR-SLC38A9 Fusion protein

## Data Availability

The data that supports the findings of this study are available in the main text and the supporting information of this article.

## Author Contributions

A.S., X.M., and T.G. were involved in the design and overall planning of the experiments. A.S. and X.M. were involved in the execution of experiments. A.S. prepared the original manuscript and X.M and T.G. edited the final manuscript.

## Acknowledgements

This study was supported by the National Institutes of Health P41GM136508 and RM1GM158451. Ampon Saeher was funded by the NCI F32CA278619. The Gonen laboratory is supported by funds from the Howard Hughes Medical Institute.

