## supplemental material for "Inhibition of the Lysosomal Amino Acid Sensor SLC38A9 by the Membrane Microprotein SPAR"

**Supplemental Materials**

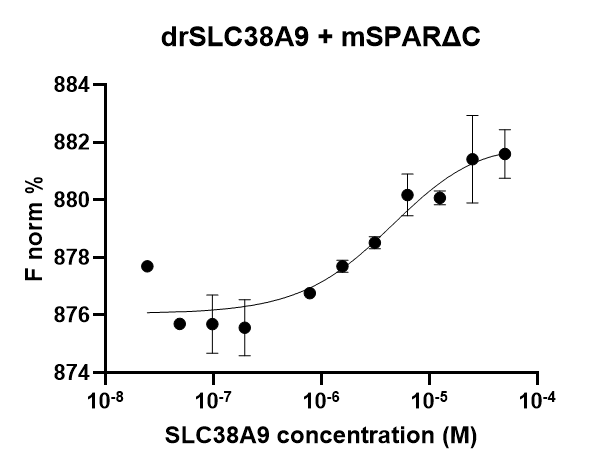

Supplemental Figure 1: MST experiment of truncated SPAR (SPAR∆C) and drSLC38A9

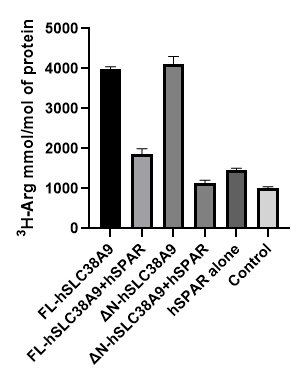

Supplemental Figure 2: ^3^H-Arg uptake experiments of MBP-cleaved hSPAR and full length human SLC38A9 (FL-hSLC38A9) and N-terminal truncated (∆N-hSLC38A9).

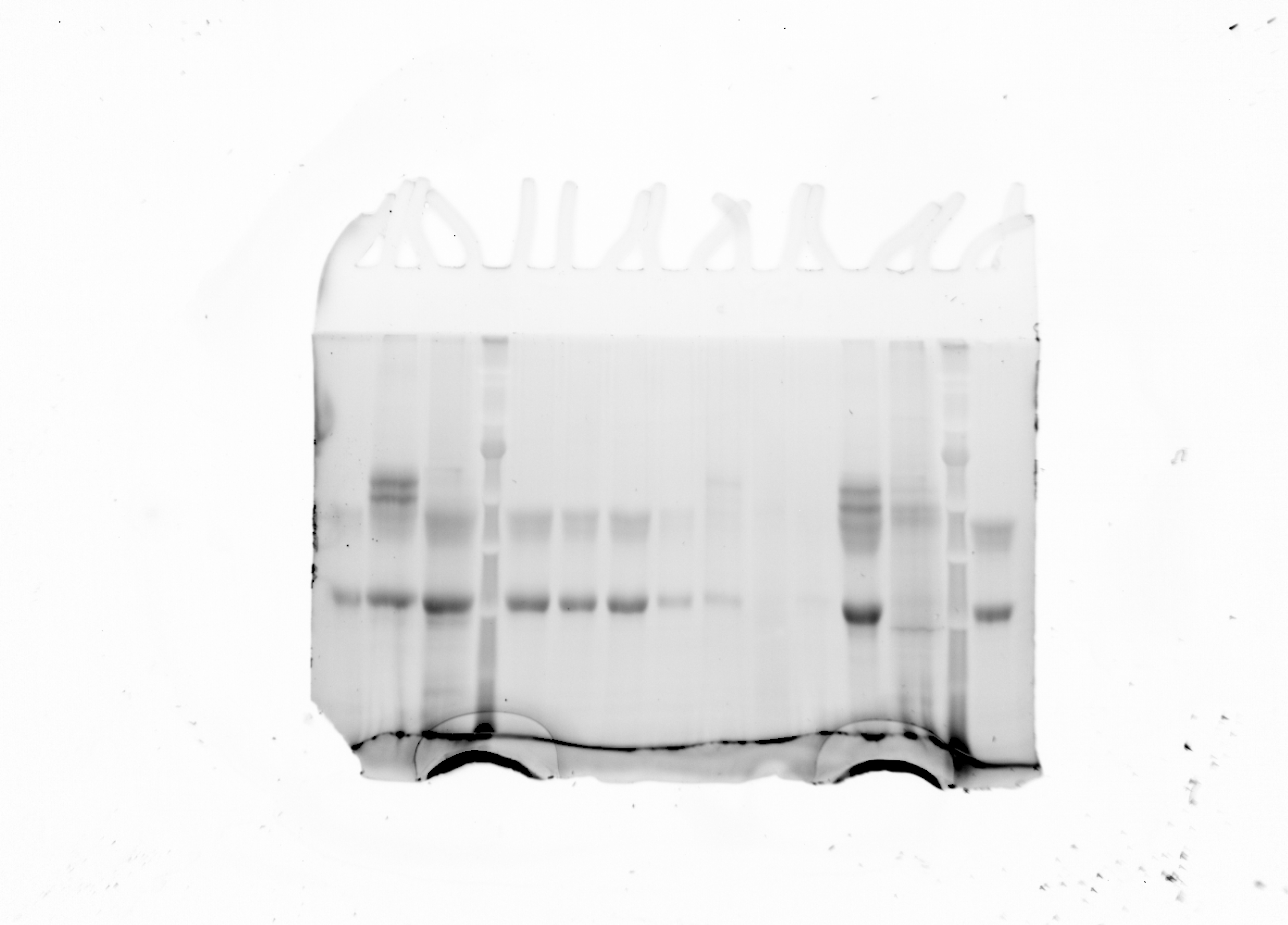

drSLC38A9

Control_3^rd^ supernatant

drSLC38A9-3^rd^ supernatant

SS-fusion_3^rd^ Supernatant

drSLC38A9_1^st^ supernatant

Control

Control-pellet

SS-fusion-pellet

SS-fusion

Control_1^st^ supernatant

drSLC38A9-pellet

SS-fusion_1^st^ supernatant

hRags proteins

Gel #1 for Anti-RagA+RagC

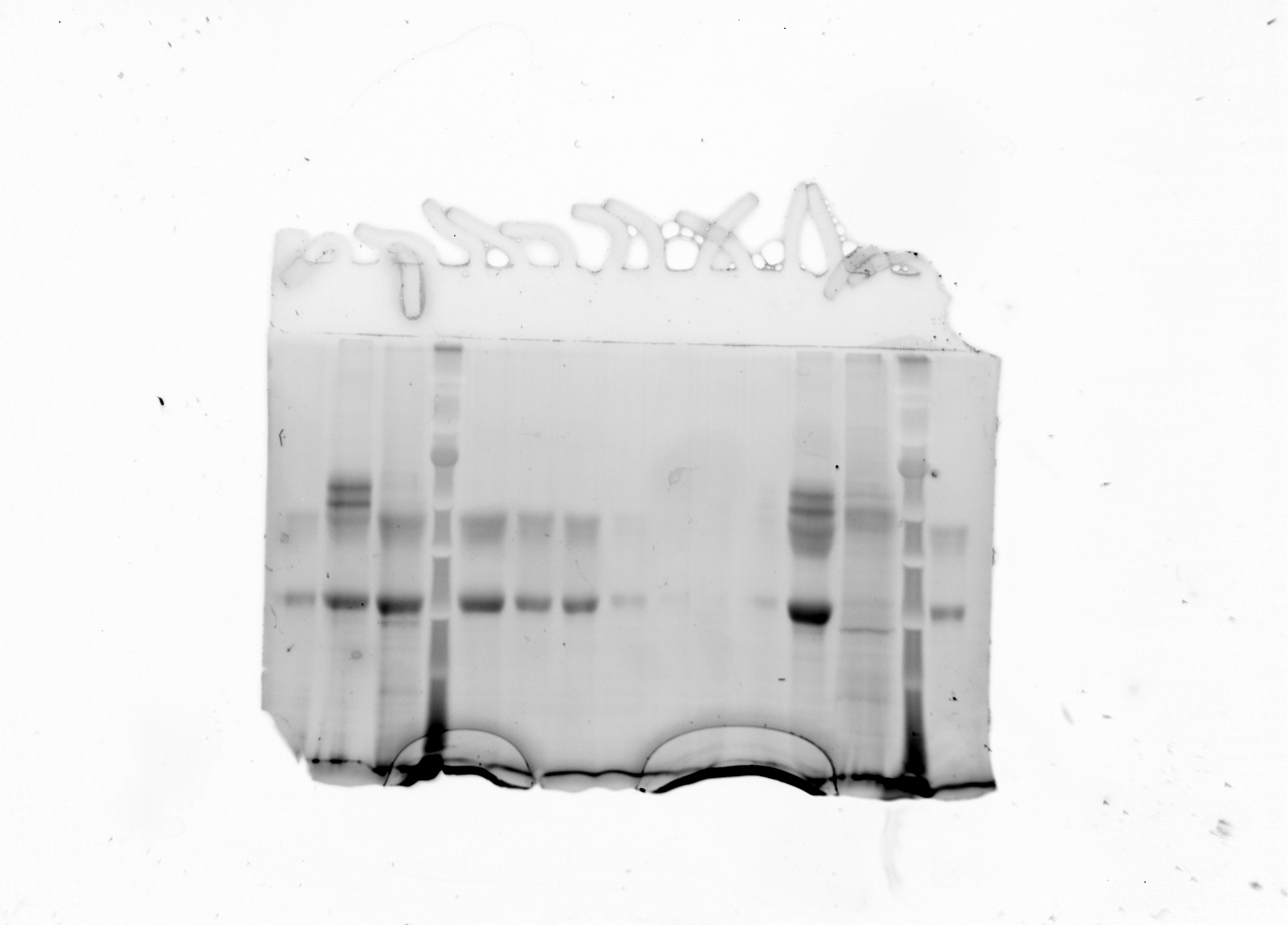

drSLC38A9

Control_3^rd^ supernatant

drSLC38A9-3^rd^ supernatant

SS-fusion_3^rd^ Supernatant

drSLC38A9_1^st^ supernatant

Control

Control

SS-fusion-pellet

SS-fusion

Control_1^st^ supernatant

drSLC38A9

SS-fusion_1^st^ supernatant

hRags proteins

Gel #2 for Anti-SLC38A9

Supplemental Figure 3: SDS PAGE gels of the proteoliposome pull down assays. These were use for western blots with anti-RagA/C and anti-SLC38A9 antibodies.

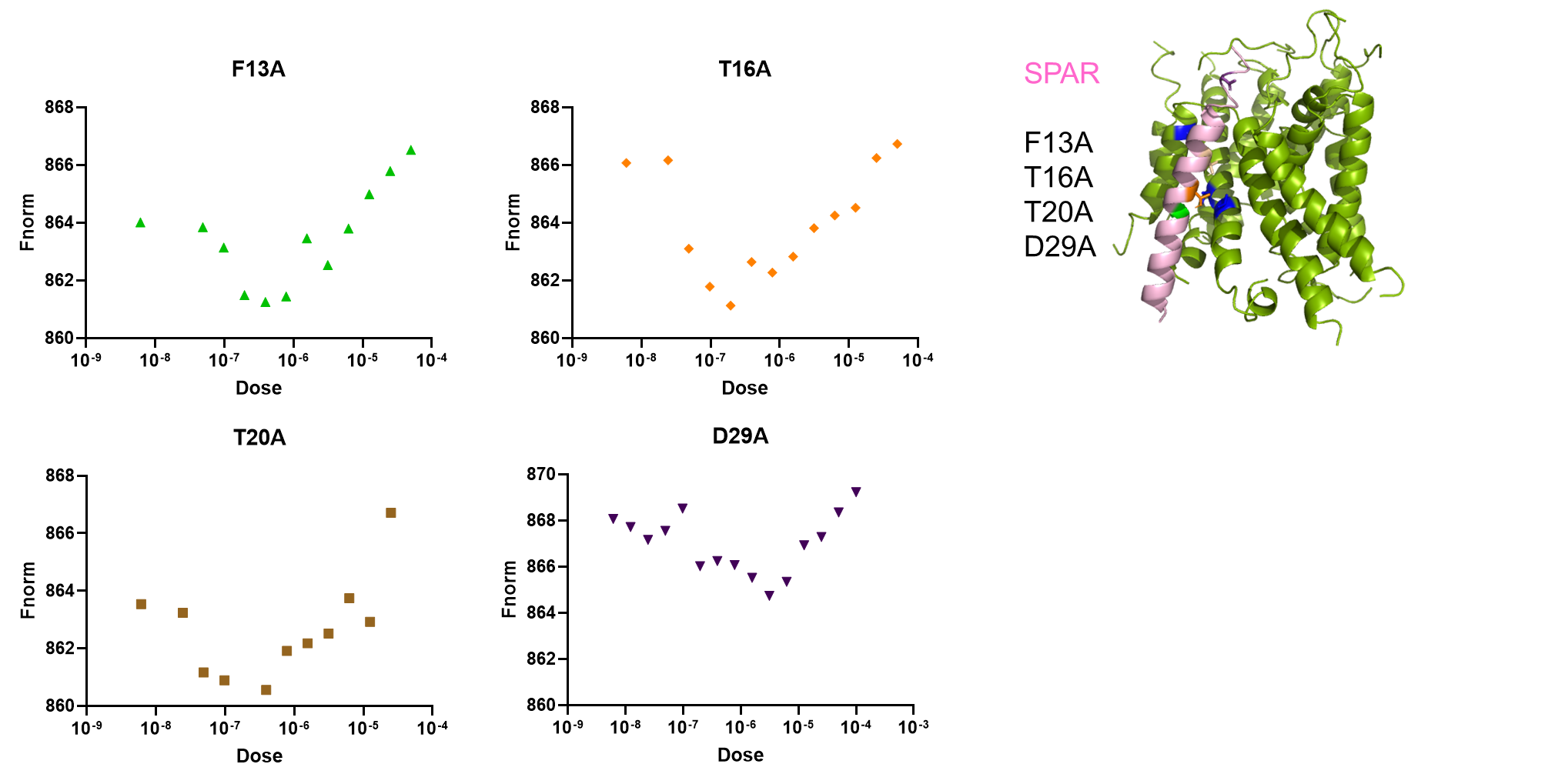

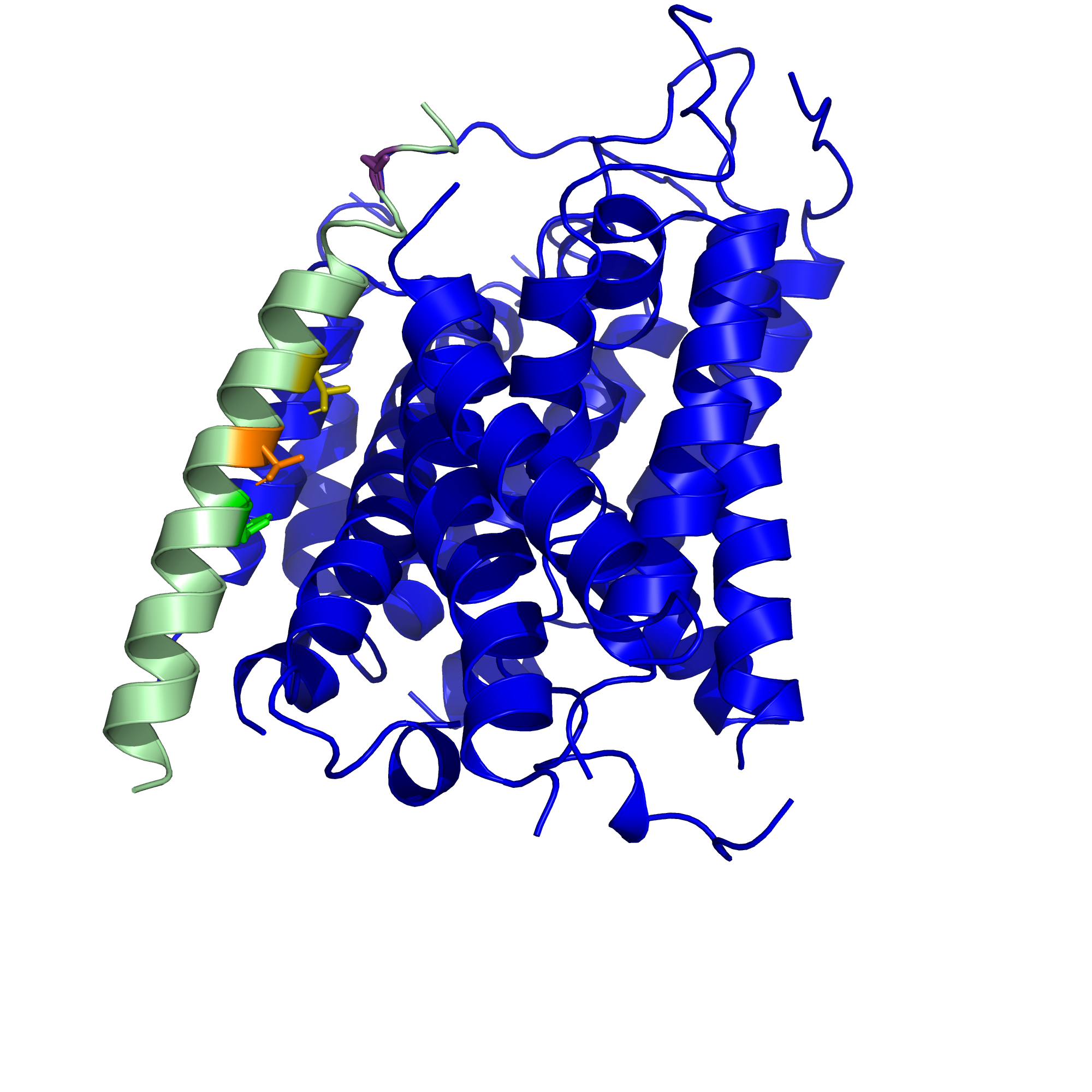

Supplemental Figure 4: Microscale Thermophoresis (MST) assays on mutated constructs of SPAR and SLC38A9

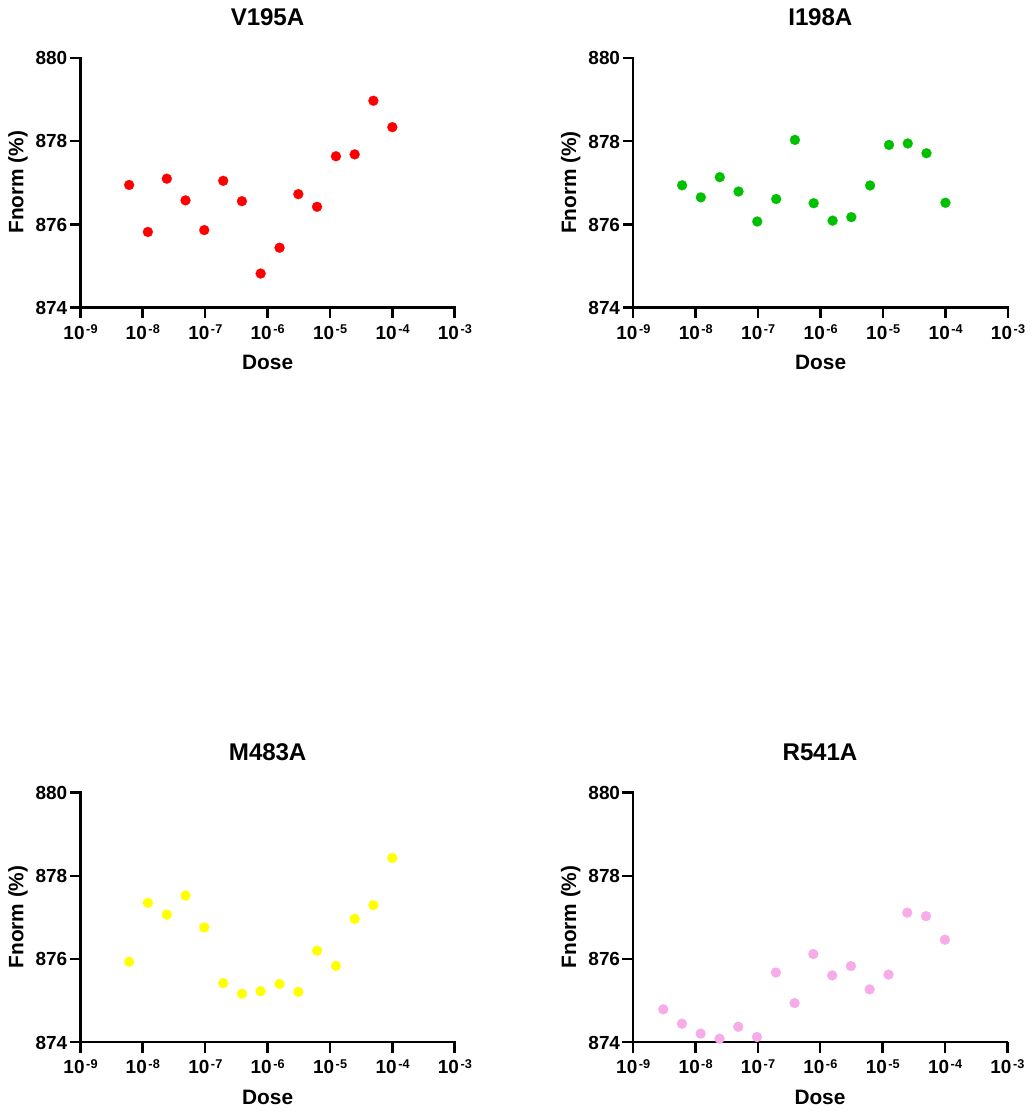

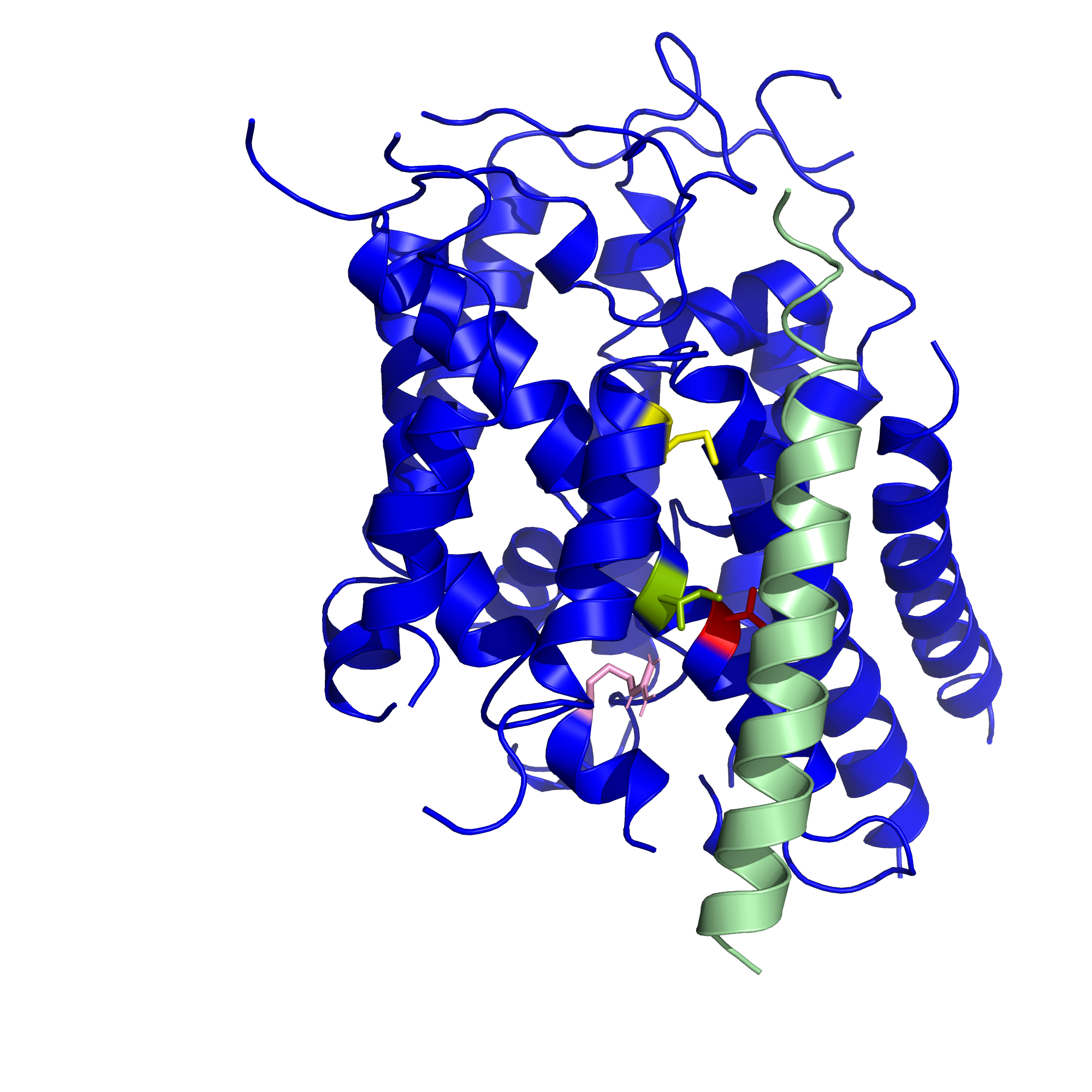

Supplemental Figure 5: Microscale Thermophoresis (MST) assays on mutated constructs of SLC38A9 and SPAR

Table 1: Constructs used for protein expressions

| Constructs | Protein Sequences |
| --- | --- |
| FL_drSLC38A9 | MHHHHHHHHHHLVPRGSMDEDSKPLLGSVPTGDYYTDSLDPKQRRPFHVEPRNIVGEDVQERVSAEAAVLSSRVHYYSRLTGSSDRLLAPPDHVIPSHEDIYIYSPLGTAFKVQGGDSPIKNPSIVTIFAIWNTMMGTSILSIPWGIKQAGFTLGIIIIVLMGLLTLYCCYRVLKSTKSIPYVDTSDWEFPDVCKYYFGGFGKWSSLVFSLVSLIGAMVVYWVLMSNFLFNTGKFIFNYVHNVNTSDAFGTNGTERVICPYPDVDPHGNSSTSLYSGSDNSTGLEFDHWWSKTNTIPFYLILLLLPLLNFRSASFFARFTFLGTISVIYLIFLVTYKAIQLGFHLEFHWFDSSMFFVPEFRTLFPQLSGVLTLAFFIHNCIITLMKNNKHQENNVRDLSLAYLLVGLTYLYVGVLIFAAFPSPPLSKECIEPNFLDNFPSSDILVFVARTFLLFQMTTVYPLLGYLVRVQLMGQIFGNHYPGFLHVFVLNVFVVGAGVLMARFYPNIGSIIRYSGALCGLALVFVLPSLIHMVSLKRRGELRWTSTLFHGFLILLGVANLLGQFFM |
| Fusion_drSLC38A9_mSPAR | MHHHHHHHHHHLVPRGSLDPKQRRPFHVEPRNIVGEDVQERVSAEAAVLSSRVHYYSRLTGSSDRLLAPPDHVIPSHEDIYIYSPLGTAFKVQGGDSPIKNPSIVTIFAIWNTMMGTSILSIPWGIKQAGFTLGIIIIVLMGLLTLYCCYRVLKSTKSIPYVDTSDWEFPDVCKYYFGGFGKWSSLVFSLVSLIGAMVVYWVLMSNFLFNTGKFIFNYVHNVQTSDAFGTQGTERVICPYPDVDPHGQSSTSLYSGSDQSTGLEFDHWWSKTNTIPFYLILLLLPLLNFRSASFFARFTFLGTISVIYLIFLVTYKAIQLGFHLEFHWFDSSMFFVPEFRTLFPQLSGVLTLAFFIHNCIITLMKNNKHQENNVRDLSLAYLLVGLTYLYVGVLIFAAFPSPPLSKECIEPNFLDNFPSSDILVFVARTFLLFQMTTVYPLLGYLVRVQLMGQIFGNHYPGFLHVFVLNVFVVGAGVLMARFYPNIGSIIRYSGALCGLALVFVLPSLIHMVSLKRRGELRWTSTLFHGFLILLGVANLLGQFFMGGGGGGMETAVIGMVAVLFVITMAITCILCYFSYDSHTQDPERSSRRSFTVATFHQEASLFTGPALQSRPLPRPQNFWTVVLVPRGSDYKDDDDK. |
| FL_hSLC38A9 | MANMNSDSRHLGTSEVDHERDPGPMNIQFEPSDLRSKRPFCIEPTNIVNVNHVIQRVSDHASAMNKRIHYYSRLTTPADKALIAPDHVVPAPEECYVYSPLGSAYKLQSYTEGYGKNTSLVTIFMIWNTMMGTSILSIPWGIKQAGFTTGMCVIILMGLLTLYCCYRVVKSRTMMFSLDTTSWEYPDVCRHYFGSFGQWSSLLFSLVSLIGAMIVYWVLMSNFLFNTGKFIFNFIHHINDTDTILSTNNSNPVICPSAGSGGHPDNSSMIFYANDTGAQQFEKWWDKSRTVPFYLVGLLLPLLNFKSPSFFSKFNILGTVSVLYLIFLVTFKAVRLGFHLEFHWFIPTEFFVPEIRFQFPQLTGVLTLAFFIHNCIITLLKNNKKQENNVRDLCIAYMLVTLTYLYIGVLVFASFPSPPLSKDCIEQNFLDNFPSSDTLSFIARIFLLFQMMTVYPLLGYLARVQLLGHIFGDIYPSIFHVLILNLIIVGAGVIMACFYPNIGGIIRYSGAACGLAFVFIYPSLIYIISLHQEERLTWPKLIFHVFIIILGVANLIVQFFMLVPRGSHHHHHHHH |
| Truncated_hSLC38A9 | MLIAPDHVVPAPEECYVYSPLGSAYKLQSYTEGYGKNTSLVTIFMIWNTMMGTSILSIPWGIKQAGFTTGMCVIILMGLLTLYCCYRVVKSRTMMFSLDTTSWEYPDVCRHYFGSFGQWSSLLFSLVSLIGAMIVYWVLMSNFLFNTGKFIFNFIHHIADTDTILSTANSNPVICPSAGSGGHPDASSMIFYAADTGAQQFEKWWDKSRTVPFYLVGLLLPLLNFKSPSFFSKFNILGTVSVLYLIFLVTFKAVRLGFHLEFHWFIPTEFFVPEIRFQFPQLTGVLTLAFFIHNCIITLLKNNKKQENNVRDLCIAYMLVTLTYLYIGVLVFASFPSPPLSKDCIEQNFLDNFPSSDTLSFIARIFLLFQMMTVYPLLGYLARVQLLGHIFGDIYPSIFHVLILNLIIVGAGVIMACFYPNIGGIIRYSGAACGLAFVFIYPSLIYIISLHQEERLTWPKLIFHVFIIILGVANLIVQFFMLVPRGSHHHHHHHH |
| Human SPAR | MKIEEGKLVIWINGDKGYNGLAEVGKKFEKDTGIKVTVEHPDKLEEKFPQVAATGDGPDIIFWAHDRFGGYAQSGLLAEITPDKAFQDKLYPFTWDAVRYNGKLIAYPIAVEALSLIYNKDLLPNPPKTWEEIPALDKELKAKGKSALMFNLQEPYFTWPLIAADGGYAFKYENGKYDIKDVGVDNAGAKAGLTFLVDLIKNKHMNADTDYSIAEAAFNKGETAMTINGPWAWSNIDTSKVNYGVTVLPTFKGQPSKPFVGVLSAGINAASPNKELAKEFLENYLLTDEGLEAVNKDKPLGAVALKSYEEELVKDPRIAATMENAQKGEIMPNIPQMSAFWYAVRTAVINAASGRQTVDEALKDAQTNSSSNNNNNNNNNNLGIEGRISEFMETAVIGVVVVLFVVTVAITCVLCCFSCDSRAQDPQGGPGRSFTVATFRQEASLFTGPVRHAQPVPSAQDFWTFMLVPRGSDYKDDDDK |
| Mouse SPAR | MKIEEGKLVIWINGDKGYNGLAEVGKKFEKDTGIKVTVEHPDKLEEKFPQVAATGDGPDIIFWAHDRFGGYAQSGLLAEITPDKAFQDKLYPFTWDAVRYNGKLIAYPIAVEALSLIYNKDLLPNPPKTWEEIPALDKELKAKGKSALMFNLQEPYFTWPLIAADGGYAFKYENGKYDIKDVGVDNAGAKAGLTFLVDLIKNKHMNADTDYSIAEAAFNKGETAMTINGPWAWSNIDTSKVNYGVTVLPTFKGQPSKPFVGVLSAGINAASPNKELAKEFLENYLLTDEGLEAVNKDKPLGAVALKSYEEELVKDPRIAATMENAQKGEIMPNIPQMSAFWYAVRTAVINAASGRQTVDEALKDAQTNSSSNNNNNNNNNNLGIEGRISEFMETAVIGMVAVLFVITMAITCILCYFSYDSHTQDPERSSRRSFTVATFHQEASLFTGPALQSRPLPRPQNFWTVVLVPRGSDYKDDDDK |
| hSPAR with cleavable Tag | MKIEEGKLVIWINGDKGYNGLAEVGKKFEKDTGIKVTVEHPDKLEEKFPQVAATGDGPDIIFWAHDRFGGYAQSGLLAEITPDKAFQDKLYPFTWDAVRYNGKLIAYPIAVEALSLIYNKDLLPNPPKTWEEIPALDKELKAKGKSALMFNLQEPYFTWPLIAADGGYAFKYENGKYDIKDVGVDNAGAKAGLTFLVDLIKNKHMNADTDYSIAEAAFNKGETAMTINGPWAWSNIDTSKVNYGVTVLPTFKGQPSKPFVGVLSAGINAASPNKELAKEFLENYLLTDEGLEAVNKDKPLGAVALKSYEEELVKDPRIAATMENAQKGEIMPNIPQMSAFWYAVRTAVINAASGRQTVDEALKDAQTNSSSNNNNNNNNNNLGIEGRISEFHHHHHHHHENLYFQSGGGGGMETAVIGVVVVLFVVTVAITCVLCCFSCDSRAQDPQGGPGRSFTVATFRQEASLFTGPVRHAQPVPSAQDFWTFMLVPRGSDYKDDDDK |

Table 2: Primers for site directed mutagenesis

| V195A-SLC38A9 | 5'- ccgatcaatgaagcgagagagaagacc -3'  5'- cttctctctcgcttcattgatcggagc -3' |
| --- | --- |
| I198A-SLC38A9 | 5'- ccatagctccggccaatgaaacg -3'  5'- cgtttcattggccggagctatgg -3' |
| R451A-SLC38A9 | 5'- ctgcacagcaaccaggtaacccaag -3'  5'- cttgggttacctggttgctgtgcag -3' |
| M483A-SLC38A9 | 5'- gaacctcgccgccaggactcct -3'  5'- caggagtcctggcggcgagg -3' |
| F13A-SPAR | 5’gttgtagtagtgttggctgttgttactgtggcgat3’  3’caacatcatcacaaccgacaacaatgacaccgcta5’ |
| T16A-SPAR | 5’gttgtagtagtgttgtttgttgttgctgtggcgat3’ 75.6C  3’caacatcatcacaacaaacaacaacgacaccgcta5’ |
| T20A-SPAR | 5’cgatcgcctgtgttctgtgctgctt3’  3’ gctagcggacacaagacacgacgaa 5’ |
| D29A-SPAR | 5’gcttcagctgcgccagccgt3’  3’cgaagtcgacgcggtcggc 5’ |
